# T cell receptor-dependent S-acylation of ZAP-70 controls activation of T cells

**DOI:** 10.1101/2020.06.30.180885

**Authors:** Ritika Tewari, Bieerkehazhi Shayahati, Ying Fan, Askar M. Akimzhanov

**Author notes:** To whom correspondence may be addressed: 6431 Fannin St., Suite 6.200, Department of Biochemistry and Molecular Biology, University of Texas-McGovern Medical School, Houston, TX, 77030. **E-mail:**.

## Abstract

ZAP-70 is a cytoplasmic tyrosine kinase essential for T cell-mediated immune responses. Upon engagement of the T cell receptor, ZAP-70 is quickly recruited to the specialized plasma membrane domains, becomes activated and released to phosphorylate its laterally segregated downstream targets. A shift in ZAP-70 distribution at the plasma membrane is recognized as a critical step in T cell receptor signal transduction and amplification. However, the molecular mechanism supporting stimulation-dependent plasma membrane compartmentalization of ZAP-70 remains poorly understood. In this study, we identified previously uncharacterized reversible lipidation (S-acylation) of ZAP-70. We found that this post-translational modification of ZAP-70 is dispensable for its enzymatic activity. However, the lipidation-deficient mutant of ZAP-70 failed to propagate the T cell receptor signaling cascade suggesting that S-acylation is essential for ZAP-70 interaction with its protein substrates. The kinetics of ZAP-70 S-acylation were consistent with early T cell signaling events indicating that agonist-induced S-acylation is a part of the signaling mechanism controlling T cell activation and function.

**Significance Statement:** Activation of T cells is a critical part of the adaptive immune response to pathogen exposure. We found that ZAP-70, a regulatory protein essential for T cell activation, can undergo a post-translational modification with long chain fatty acids, known as S-acylation. In this report, we show that S-acylation of ZAP-70 is T cell receptor-dependent and required for its signaling function. We found that loss of ZAP-70 S-acylation resulted in T cell unresponsiveness to T cell receptor stimulation indicating that abnormalities in protein S-acylation can potentially contribute to the T cell immunodeficiency disorders.

## Introduction

ZAP-70 (“CD3ζ-chain-associated protein kinase 70”) is one of the first proteins activated upon T cell receptor (TCR) engagement by the peptide/major histocompatibility complex on the surface the antigen-presenting cell (1). Stimulation of the TCR leads to activation of the Src-family kinase Lck which then phosphorylates immunoreceptor tyrosine-based activation motifs (ITAMs) of TCR-associated CD3ζ-chain (1, 2). Double phosphorylation of ITAMs creates a high-affinity docking site for Src homology 2 (SH2) domains of ZAP-70 resulting in its recruitment to the TCR-CD3 complex. Binding to ITAMs triggers conformational changes that make ZAP-70 more accessible to phosphorylation by Lck and subsequently leads to ZAP-70 autophosphorylation and full activation (3–5). Activated ZAP-70 is released from ITAMs and proceeds to target its effectors, primarily membrane-bound scaffolding proteins LAT and SLP-76 (6). Phosphorylation of LAT and SLP-76 nucleates the assembly of the multiprotein signaling complex which further propagates the downstream TCR signaling cascade resulting in activation of the phospholipase PLC-γ1, cytoplasmic calcium release from ER, and ultimately, transcriptional responses associated with T cell activation and clonal expansion (6–9).

Thus, the signaling function of the ZAP-70 kinase relies on two temporally and spatially segregated events. During the first step, the phosphorylated TCR-CD3 complex amplifies the initial antigenic signal through repeated cycles of ZAP-70 recruitment, activation and release (9). Through the next step, activated ZAP-70 translocates into distinct plasma membrane domains where it can access its protein substrates (9–11). This compartmentalization of early phosphorylation events not only ensures rapid TCR signal amplification and distribution but can also help to avoid premature or unspecific T cell activation. It is not clear, however, what mechanism prevents activated and decoupled ZAP-70 from being dispersed back into the cytoplasm and promotes its prompt translocation to the laterally segregated downstream targets.

It has been proposed that both plasma membrane anchoring and lateral distribution of proteins within the plasma membrane can be mediated by S-acylation – reversible lipidation of cysteine thiols with long-chain fatty acids (12, 13). In particular, this post-translational modification (also referred to as palmitoylation) was found to be essential for biological activity of two immediate ZAP-70 effectors, Lck and LAT (14, 15). It has been shown that while still catalytically active, the acylation-deficient mutant of Lck was unable to phosphorylate ITAMs and failed to activate ZAP-70, thus leading to abrogated TCR signaling (14, 16, 17). Furthermore, our previous study demonstrated that initiation of the Fas receptor signaling pathway in T cells relies on rapid agonist-induced changes in Lck S-acylation levels and inhibition of this modification rendered T cells completely resistant to Fas-mediated apoptosis (18). Correspondingly, S-acylation of LAT was found to be essential for its correct targeting to the TCR signaling assemblies and propagation of downstream signaling events, including phosphorylation of PLC-γ1, calcium influx and activation of transcription factors (15, 19–22).

In this study, we report previously unknown S-acylation of ZAP-70. We found that this post-translational modification is TCR-dependent and identified the cysteine in position 564 as the primary site of ZAP-70 S-acylation. Mutation of this cysteine residue did not affect the catalytic ability of the kinase but led to extended interaction of ZAP-70 with its activator kinase Lck, resulting in significantly increased ZAP-70 phosphorylation levels. However, the acylation-deficient mutant of ZAP-70 failed to phosphorylate its downstream targets, adaptors LAT and SLP-76, causing disruption of the TCR signaling chain. Overall, our study describes a previously uncharacterized post-translational modification of ZAP-70 that serves as a critical regulator of its signaling function in T cell activation.

## Results

### ZAP-70 is S-acylated at cysteine 564

We first identified ZAP-70 as an S-acylated protein in our initial experiments aimed at detection of novel lipidated proteins in the proximal TCR signaling pathway. To survey T cell protein lipidation, we employed the Acyl-Biotin Exchange (ABE) assay, a method allowing for rapid and highly sensitive detection of S-acylated proteins (23). Briefly, free thiol groups in proteins in Jurkat T cell lysate were blocked with methyl methanethiosulfonate (MMTS), followed by cleavage of thioester bonds with neutral hydroxylamine (HA). The newly exposed thiol groups were then biotinylated, pulled down using streptavidin beads and the captured proteins were analyzed by immunoblotting. This approach, in combination with a candidate-based screening of the proximal TCR signaling components, allowed us to detect previously uncharacterized S-acylation of several key TCR signaling proteins, including adaptor protein GRB2, phospholipase PLC-γ1 and tyrosine kinase ZAP-70 (**Fig. 1*A***). These findings suggested that this post-translational modification is more widespread than initially thought and prompted us to investigate a possible role of S-acylation in regulation of the ZAP-70 signaling function.

**Fig. 1.**
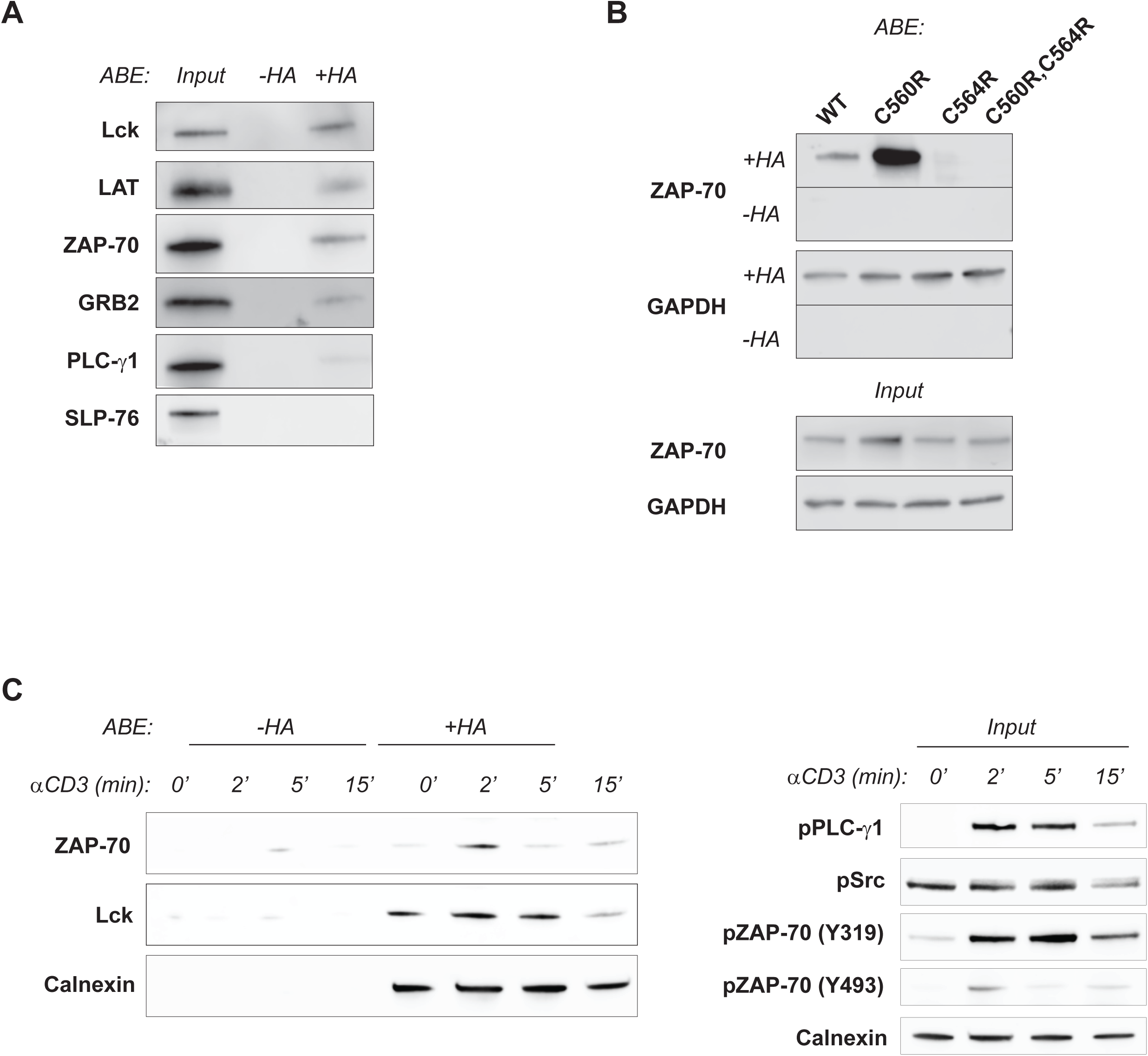
ZAP-70 is S-acylated at cysteine 564. *(A)* Identification of S-acylation targets in Jurkat T cells. S-acylated proteins were captured using ABE and detected using protein-specific antibodies. Samples not treated with hydroxylamine (-HA) were used to estimate nonspecific binding with streptavidin resin. Lck and LAT were used as the positive control. PLC-γ1, GRB2, and ZAP-70 were identified as novel S-acylated proteins. Input samples shown were collected before addition of streptavidin. *(B)* Identification of ZAP-70 S-acylation site. ABE was performed on P116 cells stably expressing WT or mutant versions of ZAP-70. Loss of ZAP-70 S-acylation was observed in cells expressing ZAP-70 with C564R mutation and ZAP-70 with C560R, C564R double-mutation. S-acylation of GAPDH, a known S-acylated protein, was not affected and served as a positive and loading control. *(C)* Kinetics of agonist-induced S-acylation. Jurkat cells were stimulated with 10 µg anti-CD3 antibody for the indicated times and lysates were subject to ABE analysis. Calnexin, a known S-acylated protein was used as a positive and loading control. Input samples were used to confirm phosphorylation of T cell signaling proteins in response to T cell stimulation.

Our next goal was to determine the site of ZAP-70 S-acylation. To identify the modified cysteine residues, we analyzed ZAP-70 protein sequence using CSS-Palm 4.0 software (24). Based on the CSS-Palm 4.0 prediction, two cysteine residues at positions 560 and 564 had an increased probability of being S-acylated. Due to the limited accessibility of C560, we hypothesized that C564 in the C-terminal part of the protein is the main site of S-acylation (**Fig. 2*A***). The hypothesis was further supported by a study reporting that a homozygous *ZAP70* mutation of this residue (c.1690T>C; p.C564R) was associated with severe combined immunodeficiency in a human patient (25). To experimentally validate S-acylation of C564, we used ZAP-70-deficient P116 Jurkat T cells to generate T cell lines stably expressing ZAP-70 mutants carrying either single or combinatorial substitutions of cysteines in positions 560 and 564. Using the ABE assay, we found that the substitution of the cysteine 564 by an arginine resulted in loss of ZAP-70 S-acylation (**Fig. 1*B***), indicating that C564 is indeed requred for this post-translational modification.

**Fig. 2.**
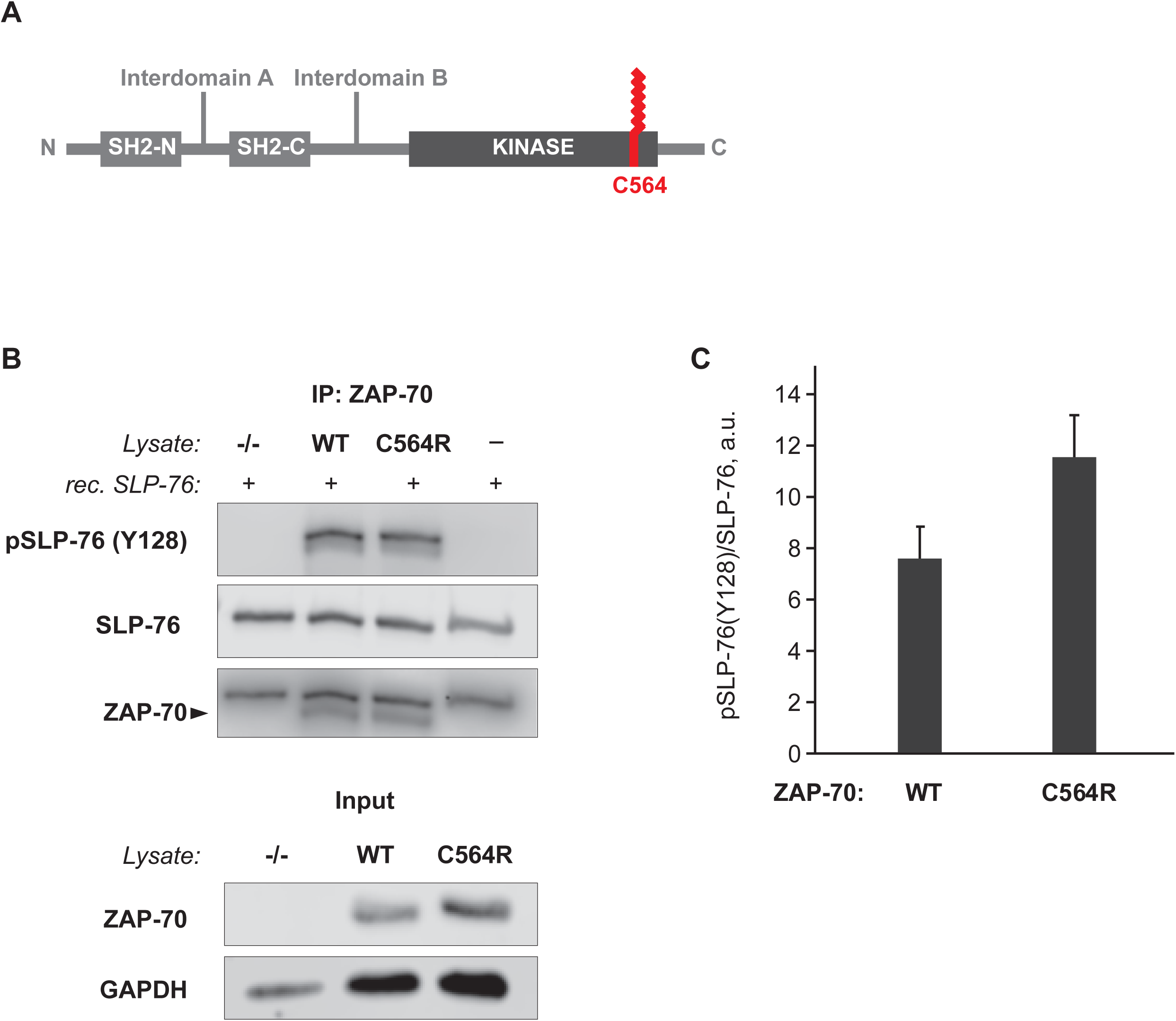
Acylation-deficient mutant of ZAP-70 is catalytically active. *(A)* Schematic representation of ZAP-70 showing the position of the S-acylated cysteine residue (C564) in the kinase domain. *(B) In vitro* kinase assay performed using ZAP-70 immunoprecipitated from P116 (ZAP-70 null) Jurkat T cells expressing WT or acylation-deficient C564R ZAP-70. Phosphorylation of SLP-76 was detected using phospho-SLP-76 (Y128) antibody. *(C)* Quantified data showing phosphorylation of SLP-76 by WT or C564R ZAP-70. Data shown are representative of three independent experiments and represented as mean ± SEM, normalized to total SLP-76.

### S-acylation of ZAP-70 is TCR-dependent

S-acylation of ZAP-70 was initially detected in resting Jurkat T cells. Considering the highly dynamic nature of early TCR signaling, we hypothesized that stimulation of the TCR can trigger very rapid changes in ZAP-70 S-acylation levels. To test this hypothesis, we treated Jurkat T cells with anti-CD3 antibody to activate the TCR signaling pathway and assessed the kinetics of ZAP-70 lipidation using the ABE assay. We found that T cell stimulation resulted in robust but transient increase in S-acylation of ZAP-70, peaking at approximately 2 minutes after TCR engagement (**Fig. 1*C***). Interestingly, another proximal signaling kinase, Lck, showed different kinetics with more sustained S-acylation lasting up to 5 minutes post-stimulation, possibly indicating distinct enzymatic control of its lipidation. Additionally, to determine whether transient increase in S-acylation of ZAP-70 depends on elevations in cytoplasmic calcium, we performed a similar experiment using J.gamma1 (PLC-γ1 null) Jurkat T cells deficient in TCR-mediated calcium release (26). We found that loss of calcium signaling did not affect basal levels of ZAP-70 S-acylation, but the TCR-induced changes in ZAP-70 lipidation were completely absent in this cell line (**Fig. S1**). Overall, we found that the transient increase in ZAP-70 S-acylation upon TCR stimulation closely matched the phosphorylation pattern of ZAP-70 and other TCR signaling proteins (**Fig. 1*C, input***) suggesting that similar to phosphorylation, dynamic protein S-acylation can be regulated by TCR engagement and subsequent calcium release.

### S-acylation of ZAP-70 is dispensable for its catalytic activity

A substitution of ZAP-70 cysteine in position 564 to arginine was associated with a case of severe combined immunodeficiency in a human patient (25), however, the molecular mechanism underlying this inborn error of immunity remained unresolved. The C564R mutation occurred within the C-terminal part of the ZAP-70 kinase domain (**Fig. 2*A***), suggesting that introduction of a positively charged amino acid residue in the vicinity of the ZAP-70 catalytic core could negatively affect its kinase function. To address this possibility, we set up an *in vitro* kinase assay to ascertain whether the C564R mutation can disrupt the ability of ZAP-70 to phosphorylate its protein substrate, SLP-76. To test the phosphorylation efficiency, ZAP-70 was immunoprecipitated from P116 cells stably expressing either wild type or C564R ZAP-70 and incubated with recombinant SLP-76 protein. While negative controls showed no detectable phosphorylation of SLP-76, we found that the C564R mutation did not cause any apparent loss of ZAP-70 kinase activity as evident from robust phosphorylation of SLP-76 at Y128 (**Fig. 2*B, C***). Thus, these data confirmed that the acylation-deficient mutant of ZAP-70 is catalytically active.

### Acylation-deficient ZAP-70 is phosphorylated by Lck at the plasma membrane

Since the acylation-deficient C564R mutant of ZAP-70 retained the ability to efficiently phosphorylate its substrate, we argued that loss of protein lipidation could still affect ZAP-70 activation by preventing its recruitment from the cytosol to the plasma membrane and subsequent interaction with Lck. To evaluate the effect of S-acylation on plasma membrane localization of ZAP-70, we utilized total internal reflection fluorescence (TIRF) microscopy. We transfected Jurkat P116 cells with mCherry-tagged ZAP-70 constructs and assessed the presence of fluorescent proteins in the TIRF plane of resting and stimulated T cells. We found that both wild type and C564R ZAP-70 demonstrated similar localization patterns by forming laterally mobile puncta at the plasma membrane (**Fig. 3*A***), suggesting that S-acylation is not required for ZAP-70 translocation from the cytosol to the plasma membrane.

**Fig. 3.**
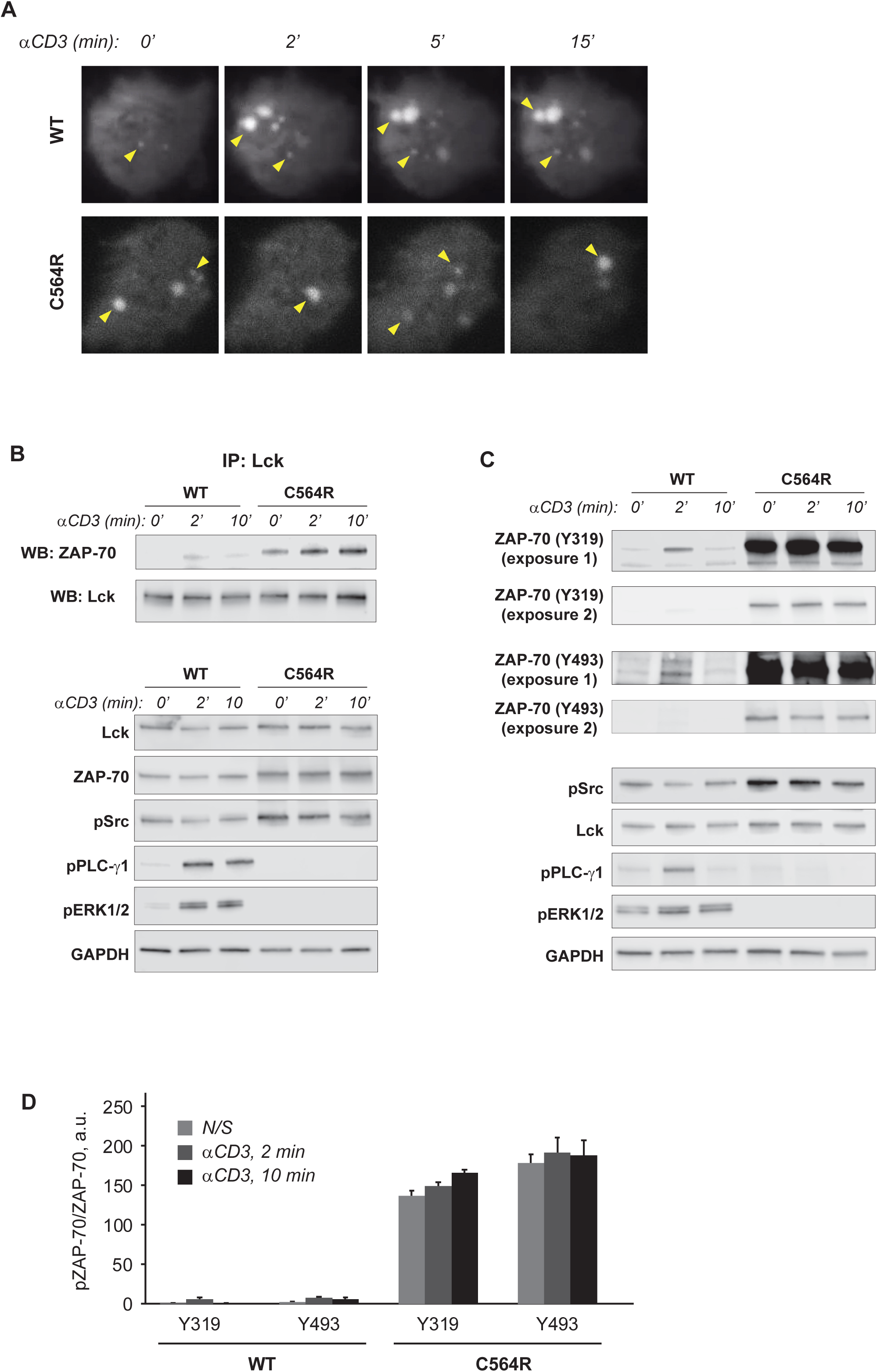
Acylation-deficient ZAP-70 exhibits increased phosphorylation at Y319 and Y493. *(A)* TIRF imaging of P116 (ZAP-70 -/-) Jurkat T cells transiently transfected with mCherry-tagged WT or acylation-deficient C564R ZAP70. Both WT ZAP-70 and C564R ZAP-70 showed similar localization patterns by forming puncta at the surface of the resting or stimulated cells. *(B)* Co-immunoprecipitation of Lck and ZAP-70. P116 Jurkat T cells stably expressing WT or C564R ZAP-70 were stimulated with anti-CD3 antibody for the indicated time points. Lck was immunoprecipitated from the lysates and the presence of ZAP-70 was assessed in eluates. *(C)* Western blot analysis of ZAP-70 phosphorylation at Y319 and Y493. P116 Jurkat T cells stably expressing WT ZAP-70 or C564R ZAP-70 were stimulated with anti-CD3 antibody for the indicated time points and phosphorylation of ZAP-70 was analyzed by immunoblotting. *(D)* Quantified data showing phosphorylation of WT or C564R ZAP-70. Data shown are representative of three independent experiments and represented as mean ± SEM, normalized to total ZAP-70.

The presence of the C564R mutant of ZAP-70 at the plasma membrane indicated that the acylation-deficient ZAP-70 can still interact with its upstream effector, Lck. To test whether S-acylation of ZAP-70 influences the interaction with its activator kinase, we immunoprecipitated Lck from P116 Jurkat cells stably expressing wild type or C564R ZAP-70. Wild type ZAP-70 became detectable in the Lck immunoprecipitate after 2 minutes of the TCR ligation by anti-CD3 antibody, but the association between these proteins rapidly decreased upon prolonged cell stimulation (**Fig. 3*B***). Surprisingly, we observed that loss of S-acylation resulted in a more sustained interaction between ZAP-70 and Lck. As shown in **Fig 3*B***, we found markedly increased co-immunoprecipitation of Lck and C564R ZAP-70 in both resting and activated cell lysates.

TCR-induced recruitment of ZAP-70 to the plasma membrane and subsequent binding to Lck facilitates phosphorylation of the ZAP-70 tyrosine residues Y319 and Y493 within the interdomain B and the tyrosine residues Y492 and Y493 in the activation loop of the catalytic domain (27). To test whether more continuous interaction of ZAP-70 with Lck affects its phosphorylation, we used anti-CD3 antibody to stimulate P116 Jurkat cells stably expressing ZAP-70 variants and assessed phosphorylation of Y319 and Y493 by immunoblotting. Consistent with more stable association of C564R ZAP-70 with Lck, phosphorylation of the acylation-deficient ZAP-70 mutant was increased by almost 250-fold compared to wild type (**Fig. 3. *C, D***). Importantly, we also observed that despite being phosphorylated at both Y319 and Y493 sites, the acylation-deficient mutant of ZAP-70 failed to propagate the TCR signaling pathway as evident from the abolished phosphorylation of proximal TCR signaling proteins PLC-γ1 and ERK1/2 (**Fig. 3*C***). These data suggest that S-acylation of ZAP-70 may could mediate its stimulus-dependent partitioning into the specific plasma membrane domains where it can interact with its downstream targets.

### S-acylation of ZAP-70 is required for initiation of TCR signaling

To further investigate the ability of C564R ZAP-70 to mediate TCR signal transduction, we stimulated P116 Jurkat cells stably rescued with ZAP-70 expression vectors with anti-CD3 antibody and examined early signaling events leading to T cell activation. As shown in **Fig. 4*A***, expression of wild type ZAP-70 rescued TCR-dependent phosphorylation of ZAP-70 substrates, LAT and SLP-76, leading to subsequent activation of downstream mediators of the proximal TCR pathway, PLC-γ1 and ERK1/2. In contrast, we found that the acylation-deficient C564R mutant of ZAP-70 failed to phosphorylate its targets, resulting in disruption of the TCR signaling cascade (**Fig. 4*A***). Notably, expression of the C564R ZAP-70 did not affect activation of upstream Src-family kinases Lck and Fyn.

**Fig. 4.**
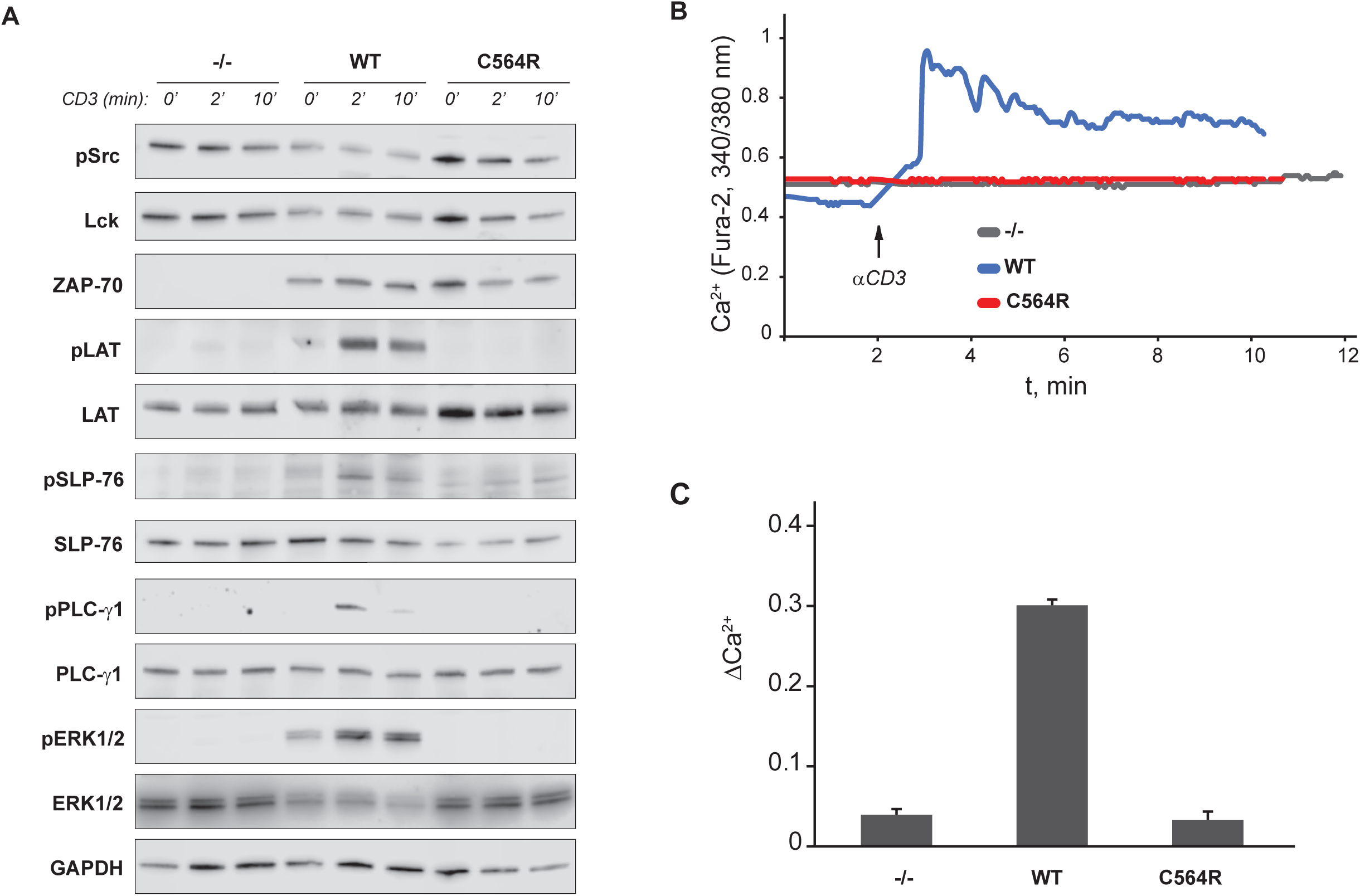
S-acylation of ZAP-70 is required for proximal TCR signaling. *(A)* Phosphorylation of proximal TCR signaling proteins in P116 -(ZAP-70 -/-) Jurkat T cells stably rescued with WT or acylation-deficient C564R ZAP-70. Cells were stimulated with anti-CD3 antibody for the indicated time points and phosphorylation of TCR signaling proteins was assayed by immunoblotting. Total protein levels shown as the loading control. *(B)* TCR-dependent calcium release in P116 cells stably rescued with WT or C564R ZAP-70. Shown are representative single-cell responses measured by Fura-2 imaging. *(C)* Quantification of peak calcium influx in response to anti-CD3 stimulation. Data shown are representative of three independent experiments and represented as mean ± SEM.

Initiation of the T cell signaling events strongly depends on PLC-γ1-mediated calcium release from the ER stores (28, 29). Consistent with compromised activation of PLC-γ1, we did not detect TCR-induced cytoplasmic calcium influx in P116 cells expressing C564R ZAP-70 (**Fig. 4 *B, C***). Based on these observations, we concluded that S-acylation of ZAP-70 is essential for activation of the proximal TCR signaling pathway.

### S-acylation of ZAP-70 is essential for T cell activation

We next aimed to determine whether disruption of the early TCR signaling events observed in P116 Jurkat cells expressing acylation-deficient ZAP-70 leads to impaired T cell activation. To test the T cell transcriptional responses, we stimulated P116 Jurkat cells stably expressing ZAP-70 variants with plate-bound anti-CD3 antibody for 24 h and used ELISA to measure secretion of interleukin-2 (IL-2), a pro-inflammatory cytokine essential for T cell activation and proliferation. We found that in contrast to wild type ZAP-70, expression of C564R ZAP-70 in P116 cells resulted in suppressed production of IL-2 in response to TCR stimulation (**Fig. 5*A***). Consonantly, our flow cytometric analysis revealed that loss of ZAP-70 S-acylation led to significantly reduced expression of CD25 and CD69 T cell activation markers on the surface of stimulated cells (**Fig. 5*B-E***). The observed deficiency in T cell activation in C564R ZAP-70 expressing cells was overridden by a combination of phorbol 12-myristate 13-acetate (PMA) and ionomycin treatment, indicating that the signaling components downstream of ZAP-70 were not affected (**Fig. S2**). Together, these findings demonstrate that S-acylation of ZAP-70 is essential for T cell activation.

**Fig. 5.**
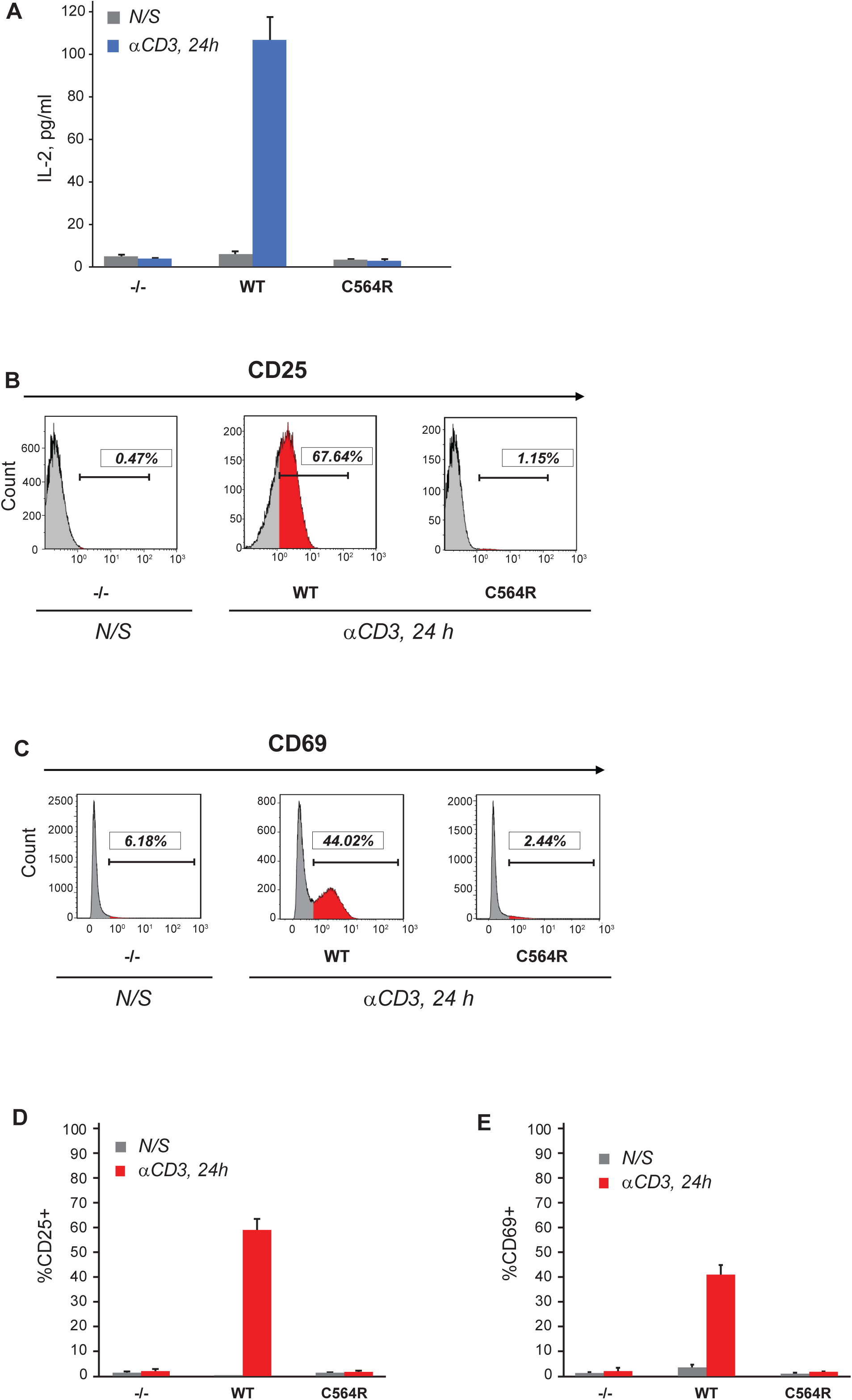
S-acylation of ZAP-70 is required for T cell activation. *(A)* IL-2 production by P116 (ZAP-70 -/-) Jurkat T cells stably rescued with WT or acylation-deficient C564R ZAP-70. IL-2 concentrations were measured by ELISA in supernatants from resting cells or cells stimulated for 24 h with plate-bound anti-CD3 antibody. Data shown are representative of three independent experiments and represented as mean ± SEM. *(B, C)* Expression of CD25 and CD69 T cell surface activation markers by P116 stably rescued with WT or C564R ZAP-70. Cells were stimulated for 24 h with plate-bound anti-CD3 antibody and analyzed by flow cytometry. *(D, E)* Quantification of CD25 and CD69 expression measured by flow cytometry. Data shown are representative of three independent experiments and represented as mean ± SEM.

## Discussion

A homozygous cysteine mutation (c.1690T>C; p.C564R) of ZAP-70 gene was reported to be associated with severe combined immunodeficiency in a human patient (24). However, the molecular mechanism underlying this inborn error of immunity remained largely unresolved. In this study, we identified C564 as the site of a post-translational modification known as protein S-acylation. Our experimental validation and functional characterization of the “disease mutant” revealed that the C564R mutation creates the acylation-deficient form of ZAP-70 that is unable to support early TCR signal transduction events resulting in inhibited T cell activation.

S-acylation of ZAP-70 was initially discovered in resting T cells; however, we found that engagement of the TCR triggers robust changes in ZAP-70 lipidation levels. We observed rapid increase in ZAP-70 S-acylation after 2 minutes of TCR stimulation followed by sharp decline to basal levels after 5 minutes (**Fig 1*C***). This rapid and transient pattern of ZAP-70 S-acylation closely resembles the phosphorylation kinetics of proximal TCR signaling proteins, including ZAP-70 itself. Thus, these observations suggest that stimulus-induced S-acylation of ZAP-70 likely plays a regulatory role during initiation of early TCR signaling events and that the enzymes controlling ZAP-70 S-acylation are an intricate part of the proximal TCR signaling machinery.

To evaluate the functional significance of ZAP-70 S-acylation, we decided to test whether the acylation-deficient mutant, C564R ZAP-70, is able to mediate activation of the TCR signaling pathway. However, we were concerned that introduction of the positively charged arginine residue could disturb the protein confirmation, resulting in ZAP-70 deficiency due to misfolding, aggregation or premature degradation. Alternatively, the C564R mutation within the C-terminal part of the kinase domain could cause a more subtle effect by affecting the ability of ZAP-70 to recognize its protein substrates. We tested both possibilities directly by stably expressing ZAP-70 constructs in P116 cells and setting up an *in vitro* kinase assay in which recombinant SLP-76 protein was added to immunoprecipitated ZAP-70 (**Fig. 2**). This experiment demonstrated that C564R ZAP-70 retained the ability to phosphorylate its substrate, indicating that this mutation did not result in protein impairment. This conclusion was further supported by our data showing that the C564R mutant can still be efficiently targeted to the plasma membrane and phosphorylated by Lck (**Fig. 3**). Moreover, substitution of the neighboring cysteine residue in the position 560 by arginine had no apparent effect on ZAP-70 signaling function as evident from the ability of C560R mutant to fully rescue T cell activation in ZAP-70 null cells (**Fig. S3**). Thus, these data suggest that the functional deficiency of the C564R mutant observed in our studies was caused by loss of S-acylation rather than diminished kinase activity.

We found that the S-acylation of ZAP-70 is critical for its signaling function in the proximal TCR pathway. Despite being catalytically active, the acylation-deficient mutant of ZAP-70 interrupted the TCR signaling cascade by failing to phosphorylate its targets, LAT and SLP-76, thus effectively blocking calcium responses and downstream signaling and subsequent T cell activation. The reason why acylation-deficient ZAP-70 was unable to phosphorylate its substrates is not entirely clear. Disproving our initial hypothesis, loss of S-acylation did not prevent the interaction between ZAP-70 and its activator kinase Lck, but rather enhanced it, resulting in significantly increased phosphorylation at Y319 and Y493 and, presumably, increased number of activated ZAP-70 molecules. A recent study published by Katz et al. showed that activated ZAP-70 is produced by Lck at the TCR-CD3 complex during multiple “catch-and-release” events that lead to amplification of the initial TCR signal (9). Upon release from the TCR, the activated ZAP-70 kinase is retained at the plasma membrane and translocates into the spatially segregated domains containing its substrates (9–11). S-acylation has been proposed to serve as a molecular basis for protein lateral mobility and sequestration into plasma membrane domains (30). Therefore, dynamic TCR-induced S-acylation can act as a mechanism supporting transient membrane anchorage of activated ZAP-70 and its subsequent migration to LAT and SLP-76-containing compartments. In this scenario, the acylation-deficient ZAP-70 mutant is continuously recruited and phosphorylated at the TCR complex but lack of lipidation prevents it from reaching its substrates and negative regulators. Future imaging studies that can afford measurement of protein distribution with nanometer precision should investigate whether S-acylation regulates ZAP-70 lateral mobility and co-localization with its substrates in stimulated T cells.

Another important question raised by our findings is the identity of enzymes responsible for mediating ZAP-70 S-acylation and deacylation upon T cell stimulation. In mammalian cells, protein S-acylation is catalyzed by a family of DHHC protein acyltransferases (PATs) (31, 32). Due to their recent identification, little is known about the role of DHHC PATs in regulation of the immune responses. Previously, we identified DHHC21 as a PAT controlling T cell activation and differentiation of naïve CD4^+^ T cells into T helper (Th) Th1 and Th2 subtypes (33). Downregulation of DHHC21 prevented S-acylation of Lck, Fyn and LAT, suggesting that this enzyme could be involved in regulation of proximal TCR signaling. However, the substrate preferences of DHHC21 towards ZAP-70 have yet to be established. Thioesterases, enzymes catalyzing cleavage of the thioester bond between the fatty acid chain and the cysteine residue, remain even more enigmatic. For a long time acyl protein thioesterases 1 and 2 (APT1, APT2) were the only enzymes known to mediate deacylation of cytosolic proteins (34, 35). Recently, the list was expanded to include a small family of ABHD17 serine hydrolases (36, 37). However, the search for other thioesterases continues and the possible role of these enzymes in regulation of the immune system remains to be determined.

In conclusion, our study describes a previously uncharacterized post-translational modification of ZAP-70 with long chain fatty acids, known as S-acylation. Our biochemical and functional analysis of this modification established stimulus-dependent S-acylation of ZAP-70 as a novel signal transduction mechanism critically supporting the physiological function of ZAP-70 in T cell-mediated immunity.

## Materials and Methods

### Antibodies and reagents

Antibodies against the indicated proteins were purchased from Cell Signaling Technology: Lck (Cat. 2752), pSrc (Cat. 2101), pZAP70 (Y319, Cat. 2717), pZAP-70 (Y493, Cat. 2704), ZAP-70 (Cat. 3165), pErk1/2 (Cat. 9101), Erk1/2 (Cat. 4695), GAPDH (Cat. 5174), Calnexin (Cat. 2679), LAT (Cat. 9166), pLAT (Cat. 3581), PLC-γ1 (Cat. 2822), pPLCγ1 (Cat. 2821), SLP-76 (Cat. 70896) and GRB2 (Cat. 3972). Antibodies against pSLP-76 were purchased from Invitrogen, Cat. PA5-39759. Following antibodies were used for immunoprecipitation-anti-Lck clone 3A5 (EMD Millipore, Cat. 05-435) and anti-ZAP-70 clone 35A.2 (EMD Millipore, Cat. 05-253). Following reagents were used-Anti-human CD3 antibody (OKT3) (eBioscience™, Cat. 14-0037-82), Pierce™ Protein A agarose beads (Thermo Scientifc™, Cat. 20333), Hydroxylamine hydrochloride (Sigma, Cat.159417), Methyl methanethiosulfonate (MMTS) (Sigma, Cat. 208795), n-Dodecyl β-D-maltoside (DDM) (Sigma, Cat. D4641), EZ-Link™ HPDP-biotin (Thermo Scientific™, Cat. 21341), Streptavidin agarose beads (Invitrogen, S951), Poly-L-lysine (Sigma, Cat. P4707), Phosphatase Inhibitor Cocktail 2 (Sigma, Cat. P5726), cOmplete Protease Inhibitor Cocktail tablets (Roche, Cat. 11836170001) and ML211 (Cayman Chemical, Cat. 17630).

### Cell culture

Jurkat E6-1 (Jurkat clone E6-1 (ATCC® TIB-152™), P116 (ATCC® CRL-2676™) and P116-cl39 (ATCC® CRL-2677™) cells were purchased from American Type Culture Collection (ATCC). Cells were maintained in RPMI 1640 medium (Corning) with 2 mM L-glutamine adjusted to contain 4.5 g/l glucose, 1.0 mM sodium pyruvate, 1% penicillin/streptomycin and 10% fetal bovine serum. Stably transfected cell lines were cultured in complete medium containing 0.5 μg/ml puromycin.

### DNA constructs

hZAP-70 was cloned into pLEX_307 vector (Addgene, #41392) using Gateway cloning strategy (https://www.thermofisher.com/us/en/home/life-science/cloning/gateway-cloning/gateway-technology.html). Site-directed mutagenesis was performed by GenScript to generate C564R ZAP-70, C560R ZAP-70 and C560R, C564R ZAP-70 in the pLEX_307 vector backbone. WT ZAP-70 or C564R ZAP-70 was cloned into the pmCherry-C1 vector from Clontech (now Takara Bio; 632524; Mountain View, CA) using the NEBuilder high-fidelity DNA assembly cloning (NEB; E5520) with the following forward primer-cgacggtaccgcgggcccgggatccATGCCAGACCCCGCGGCG and reverse primer-tcagttatctagatccggtggatcCTACGTAGAATCGAGACCGAGGAGAGGGTTAG as per the manufacturer’s protocol.

### Western blotting

Protein samples in SDS sample buffer (50 mM Tris-HCl (pH 6.8), 2 % SDS, 10 % glycerol, 5 % β-mercaptoethanol and 0.01 % bromophenol blue) were incubated at 100 °C for 5 minutes, resolved by SDS-PAGE and transferred to nitrocellulose membrane. After transfer, membranes were blocked with 5 % bovine serum albumin (BSA) in PBS-T (0.1 % Tween-20 in PBS buffer) and incubated with primary antibodies (1:1000) overnight at 4 °C followed by three PBS-T washes. Membranes were then incubated with HRP or Alexa-fluor conjugated secondary antibodies (1:10,000) in PBS-T for 1 hour at RT. After three washes in PBS-T membranes were imaged on the LI-COR Odyssey Scanner (LI-COR Biosciences; Lincoln, NE). Brightness and contrast were adjusted in the linear range using the Image Studio software (LI-COR).

### Lentivirus Transduction

Lentiviruses encoding WT ZAP-70, C564R ZAP-70, C560R ZAP-70 or C560R,C564R ZAP-70 were generated by transient transfection of the lentiviral vector pLEX_307 (1.2 µg) with lentiviral production plasmids, 0.6 µg of psPAX2 (Addgene, #12260) and 0.6 µg of pMD2.G (Addgene, #12259), into newly thawed HEK293T cells plated at 70% confluency in six well plates (protocol modified from Torchia et al, 2018 (38)). The supernatant containing the lentivirus was collected, spun down and aliquots were stored at −80 °C. For infection, 1 ml of 1 × 10^6^ P116 Jurkat cells was plated in twelve well plates. 1 ml of virus-containing supernatant was added to the wells at 1:1 v/v and the plate was spun at 900 × g at 30 °C for 1 h. Transduced cells were selected in puromycin containing media (0.5 μg/ml).

### Acyl-Biotin Exchange (ABE) Assay

Protein S-acylation was determined using ABE assay modified from previous publications (39– 41). We used 1 × 10^7^ cells for each condition. Cells were lysed in 600 µL of 1 % DDM lysis buffer (1% Dodecyl β-D-maltoside (DDM) in DPBS; 10 µM ML211; Phosphatase Inhibitor Cocktail 2 (1:100); Protease Inhibitor Cocktail (1 X) and PMSF (10 mM)). Post-nuclear cell lysate was subjected to chloroform-methanol (CM) precipitation by adding methanol (MeOH) and chloroform (CHCl_3_) to the lysate at a final ratio of lysate: MeOH: CHCl_3_ of 2:2:1 with vigorous shaking to create a homogeneous suspension. This was followed by spinning the tube at 10,000 x g for 5 min to form a pellet at the interphase between aqueous and organic phases. Excess solvent was aspirated and the protein pellet was resuspended in 200 µL of 2SHB buffer (2% SDS; 5 mM EDTA; 100 mM HEPES; pH 7.4) by vortexing at 42°C/1500 rpm in a thermal shaker until the pellet dissolved. Next, free thiol groups were blocked by adding 200 µL of 0.2% MMTS (S-Methyl methanethiosulfonate) in 2SHB buffer to each tube to a final concentration of 0.1% MMTS. Blocking was performed for 15 min at 42 °C with shaking at 1500 rpm in a thermal shaker. MMTS was removed by three rounds of CM precipitation and protein pellets were dissolved in 2SHB buffer and Buffer A (5 mM EDTA; 100 mM HEPES; pH 7.4) as described (41). Following three rounds of CM precipitation, 10% of the sample was retained as input control and the remaining was split into two equal volumes labeled as +HA and -HA. The +HA group was incubated with 400 mM hydroxylamine (HA, pH 7-7.2, freshly prepared) to promote thioester bond cleavage. Whereas in the -HA group, 400 mM sodium chloride was added instead of HA. Both groups were then incubated with 1 mM HPDP-biotin and 30 µl of streptavidin agarose beads overnight at RT with gentle mixing. This was followed by four rounds of washes with 1% SDS in Buffer A and the bound proteins were eluted off the beads by incubation with 4 X Laemmli SDS sample buffer supplemented with 5 mM DTT for 15 min at 80 °C with shaking. The supernatant was transferred to new tubes and 20 µL of the sample were loaded onto a 4-20 % gradient SDS-PAGE gel (Bio-Rad) and analyzed by immunoblotting.

### Immunoprecipitation

Cells were lysed in 1 % DDM lysis buffer (1% Dodecyl β-D-maltoside (DDM) in DPBS; 10 µM ML211; Phosphatase Inhibitor Cocktail 2 (1:100); Protease Inhibitor Cocktail (1 X) and PMSF (10 mM). 500 µg of lysate was used for each immunoprecipitation. 1/10 of each sample was retained as input control. For immunoprecipitation, lysates were incubated with 1 µg of anti-Lck or anti-ZAP-70 antibody and rotated overnight at 4 °C. This was followed by incubation with 30 µl of Protein A agarose beads for 4 hours at 4 °C. Interacting proteins were eluted off beads by incubation with 4 X Laemmli SDS sample buffer supplemented with 5 mM DTT for 15 min at 80 °C with shaking. Eluted proteins were analyzed by immunoblotting.

### Flow cytometry

Cell surface markers were evaluated by flow cytometry using a standardized protocol. Cells were kept on ice during all the procedures. Cells were stained with anti-human CD69 FITC (eBioscience™, Cat. 11-0699-42) or anti-human CD25 PE (eBioscience™, Cat. 12-0259-80). Detection of cell surface markers was conducted using a Beckman-Coulter Gallios Flow Cytometer (BD Biosciences, San Jose, CA, United States) and data were analyzed by Kaluza Analysis Software. Live/dead assays were determined using the Aqua Dead Cell Stain Kit (ThermoFisher, Cat. L34957).

### ELISA

0.5 × 10^6^ T cells were stimulated in 24 well tissue culture plates coated with 5 µg/ml anti-CD3 antibody for 24 h or 1 ug/ml of PMA/Ionomycin for 6 h. Following the stimulation, supernatants were collected for analysis. IL-2 concentration was measured using a human IL-2 ELISA kit (R&D Systems, DY202-05) following the manufacturer’s instructions.

### Fura-2 Calcium Imaging

Cells were loaded with Fura-2 AM as described previously (29) and placed in an imaging chamber with a glass bottom pre-coated with poly-L-lysine. Images were taken on a Nikon TiS inverted microscope (Tokyo, Japan) with a 40 X oil immersion objective. Images were taken every 3 seconds with a Photometrics Evolve electron-multiplying charged-coupled device camera (Tucson, AZ). Cells were imaged for 2 min to generate a baseline recording and then imaging media was carefully replaced with media containing 10 µg/ml of anti-CD3 antibody to initiate TCR signaling and imaged for 10 min.

### Electroporation and TIRF imaging

0.5 × 10^6^ P116 Jurkat cells were plated in six well plates. The cells were then electroporated with 10 µg mCherry-WT ZAP-70 or mCherry-C564R ZAP-70 vector at 2 pulses of 1400 V pulse voltage and 10 ms pulse width using a Neon® transfection system (Invitrogen) following manufacturer’s guidelines. Cells were incubated in antibiotic free media for 48 hours and then resuspended in imaging media (12.5 mL 4x Ion Safe solution (427.8 mM NaCl; 80 mM HEPES; 10 mM MgCl_2_; 29 mM KCl and 46 mM glucose); 5 mL 10 % BSA in water; 50 µl 1 M CaCl_2_ and water up to 50 ml) and placed in an imaging chamber with a glass bottom pre-coated with Poly-L-lysine. Images were taken on a Nikon Eclipse TiS TIRF microscope with a 60 X objective every 500 ms with an Andor Zyla scientific complementary metal oxide semiconductor camera (Belfast, Ireland). Fluorescently labeled cells were located and imaged for 3 min before treating with anti-CD3 antibody (10 μg/ml). Cells were continuously imaged for 30 min after stimulation.

### In vitro Kinase Assay

5 × 10^6^ P116 cells or P116 cells stably expressing WT ZAP-70 or C564R ZAP-70 were lysed in 500 µl of 1% DDM lysis buffer (1 % Dodecyl β-D-maltoside (DDM) in DPBS; 10 µM ML211; Phosphatase Inhibitor Cocktail 2 (1:100); Protease Inhibitor Cocktail (1 X) and PMSF (10 mM). 1/10 of each sample was retained as input control. For ZAP-70 immunoprecipitation, lysates were incubated with 1 µg of anti-ZAP-70 antibody and rotated overnight at 4 °C. This was followed by incubation with 30 µl of Protein A beads for 4 hours at 4 °C. The beads were then washed twice with the lysis buffer and twice with the kinase buffer (20 mM HEPES pH 7.5; 0.1 M NaCl; 0.05 % NP-40 and 10 mM MgCl_2_) and resuspended in 100 µl of the kinase buffer. 100 µM ATP, 1 mM DTT and 0.25 µg recombinant SLP-76 (OriGene, Cat. TP721201) were then added to the sample and incubated at 30 °C for 60 min. 4 X Laemmli SDS buffer was added to terminate the assay. The supernatant was assayed for phosphorylation of SLP-76 by immunoblotting. 0.25 µg SLP-76 alone with 100 µM ATP and 1 mM DTT in kinase buffer was used as a negative control.

## Acknowledgements

We thank Darren Boehning (Cooper Medical School of Rowan University), Debjani Dutta (National Cancer Institute, National Institutes of Health) and Ashabari Sprenger (McGovern Medical School, University of Texas Health Science Center at Houston) for helpful suggestions and shared protocols. We thank Autumn N. Marsden (McGovern Medical School, University of Texas Health Science Center at Houston) and C. Anthony Scott (Baylor College of Medicine) for technical assistance with TIRF microscopy and data analysis. We thank Savannah West (McGovern Medical School, University of Texas Health Science Center at Houston) for critical reading of the manuscript. This work was supported by National Institutes of Health Grant 1R01GM115446 and startup funds provided by the University of Texas Health Science Center at Houston (to A.M.A.).

## Supplementary figures

**Fig. S1.**
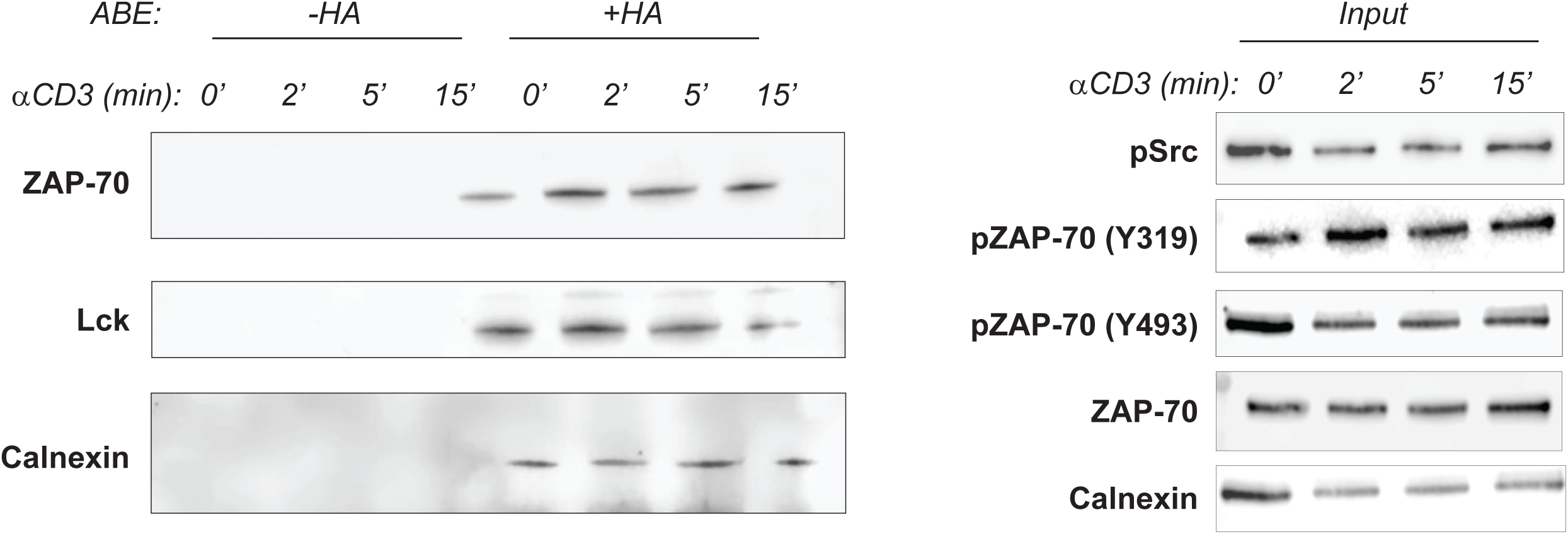
Calcium is required for TCR-induced S-acylation of ZAP-70. Kinetics of agonist-induced S-acylation assessed in J.gamma1 (PLC-γ1 null) Jurkat T cells. Cells were stimulated with 10 µg/ml anti-CD3 antibody for the indicated times and lysates were subject to ABE analysis. Calnexin, a known S-acylated protein was used as a positive and loading control. Input samples were used to confirm phosphorylation of T cell signaling proteins in response to T cell stimulation.

**Fig. S2.**
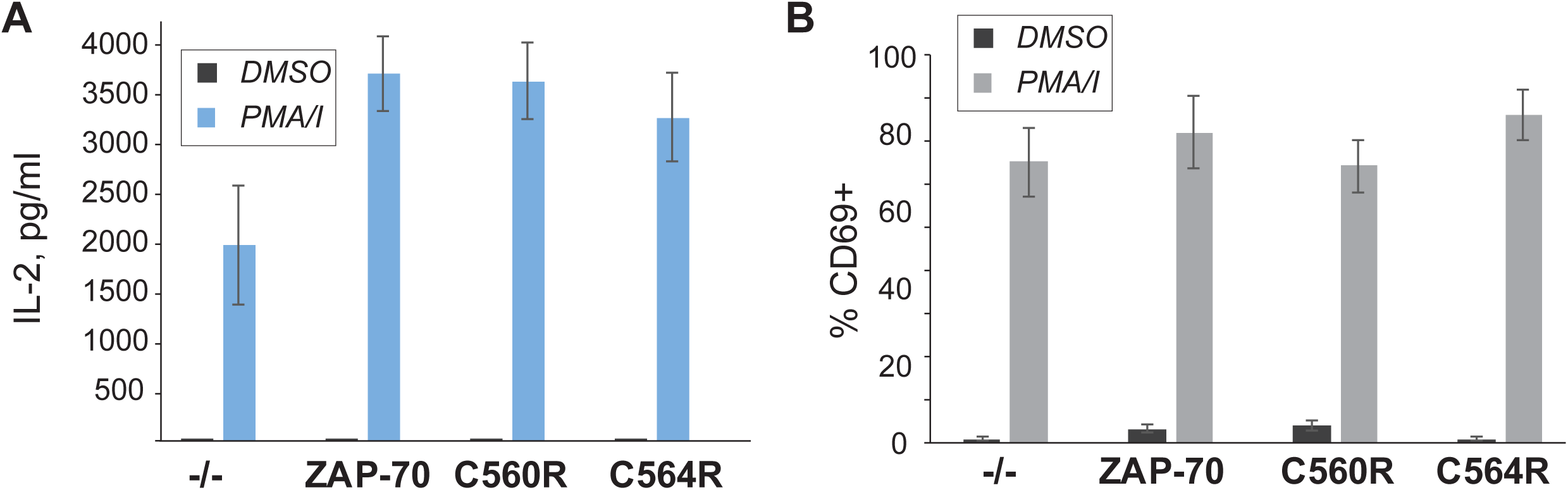
Signaling downstream of ZAP-70 is not affected in cells expressing C564R ZAP-70. *(A)* IL-2 production by P116 (ZAP-70 -/-) Jurkat T cells stably rescued with ZAP-70 variants. IL-2 concentrations were measured by ELISA in supernatants from resting cells or cells stimulated for 6 h with PMA/Ionomycin. Data shown are representative of three independent experiments and represented as mean ± SEM. *(B)* Expression of CD69 T cell surface activation markers by P116 stably rescued with ZAP-70 variants. Cells were stimulated for 6 h with PMA/Ionomycin and analyzed by flow cytometry. Data shown are representative of three independent experiments and represented as mean ± SEM.

**Fig. S3.**
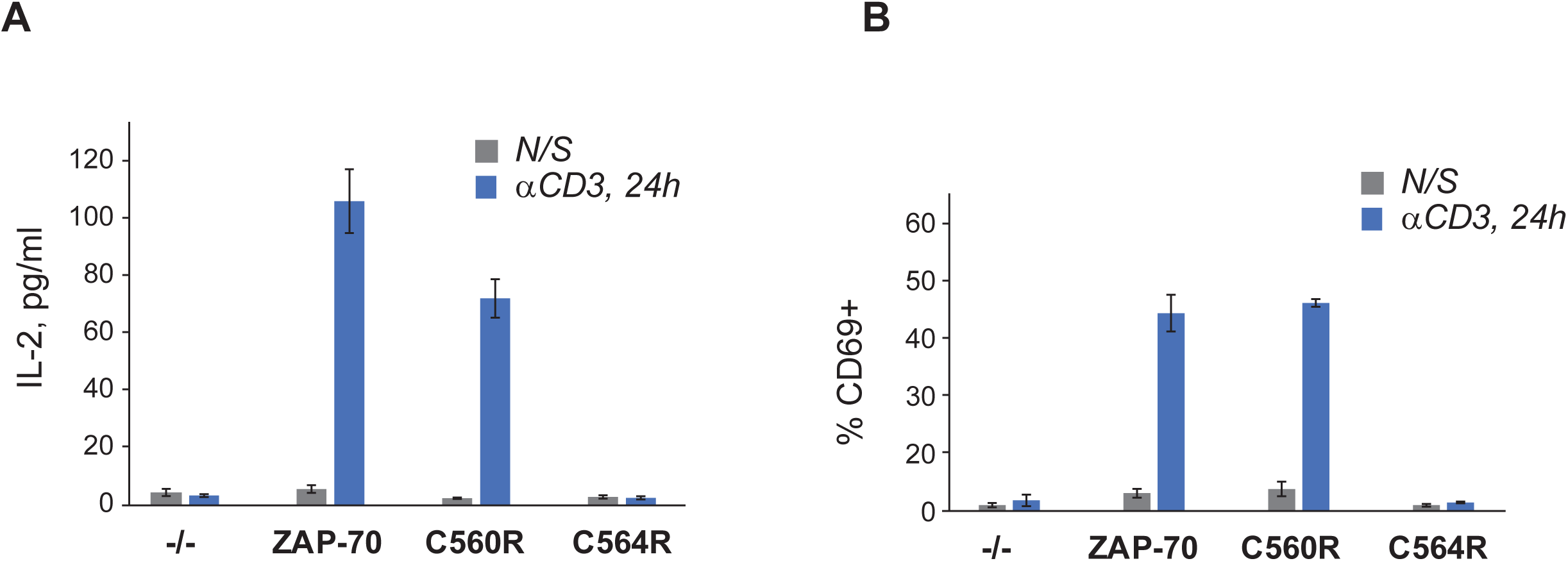
Cysteine to arginine substitution at the position 560 does not affect ZAP-70-mediated T cell activation. *(A)* IL-2 production by P116 (ZAP-70 null) Jurkat T cells stably rescued with ZAP-70 variants. IL-2 concentrations were measured by ELISA in supernatants from resting cells or cells stimulated for 24 h with plate-bound anti-CD3 antibody. Data shown are representative of three independent experiments and represented as mean ± SEM. *(B)* Expression of CD69 T cell surface activation markers by P116 stably rescued with ZAP-70 variants. Cells were stimulated for 24 h with plate-bound anti-CD3 antibody and analyzed by flow cytometry. Data shown are representative of three independent experiments and represented as mean ± SEM.

